# Bacterioplankton dynamics driven by inter-annual variation in phytoplankton spring bloom communities in the Baltic Sea

**DOI:** 10.1101/513606

**Authors:** María Teresa Camarena-Gómez, Clara Ruiz-González, Jonna Piiparinen, Tobias Lipsewers, Cristina Sobrino, Ramiro Logares, Kristian Spilling

## Abstract

In the Baltic Sea, climate change has caused shifts in the phytoplankton spring bloom communities with co-occurrence of diatoms and dinoflagellates. Such changes likely affect the composition and function of associated bacterioplankton, key members of the carbon cycling, although the actual effects are unknown. To understand how changes in phytoplankton impact on bacterioplankton composition and function, we analysed bacterioplankton communities and their production during different phases of the spring bloom in four consecutive years across the Baltic Sea, and related them to environmental variables. Phytoplankton communities varied largely in composition, modifying the taxonomic structure and richness of the associated bacterioplankton assemblages. In presence of certain diatoms *(Achnanthes taeniata, Skeletonema costatum* and *Chaetoceros* spp.), bacterial production and diversity were high and with more relative abundance of Flavobacteriia, Gammaproteobacteria and Betaproteobacteria. This bacterial community structure correlated positively with high diatom biomass and with high bacterial production rates. In contrast, during dinoflagellate-dominated blooms or when the diatom *Thalassiosira baltica* was abundant, both bacterial production rates and diversity were low, with bacterial communities dominated by SAR11 and Rhodobacteraceae. Our results demonstrate that, changes in the phytoplankton spring bloom will have profound consequences on bacterial community structure and their role in carbon cycling.

## Introduction

The Baltic Sea is suffering from a progressive increase in surface water temperature, which has been suggested to alter the food web structure, favoring small flagellates and dinoflagellates ^1^. During the last decades, shifting phytoplankton spring bloom communities from diatom-dominated blooms towards higher abundances of dinoflagellates have been reported in some subbasins of the Baltic Sea^2–4^, likely due to the increase in frequency of milder winters^1,5^ However, the consequences of these changes in spring blooms on the structure and functioning of the associated bacterioplankton communities are still poorly understood.

The nature and the timing of the spring bloom in the Baltic Sea vary between sub-basins ^6^. The bloom typically starts with the light promoting net primary production in the southernmost Baltic Sea in February/March, reaching the Gulf of Finland in April, and the Gulf of Bothnia in May (Map in Fig. 1). The bloom reaches the peak at the time when inorganic nutrients have been depleted; N-limitation prevails in most of the Baltic Sea except for the P-limited Bay of Bothnia^7–9^. The subsequent decline phase is characterized by rapid sinking of the phytoplankton cells ^10^.

**Figure 1.**
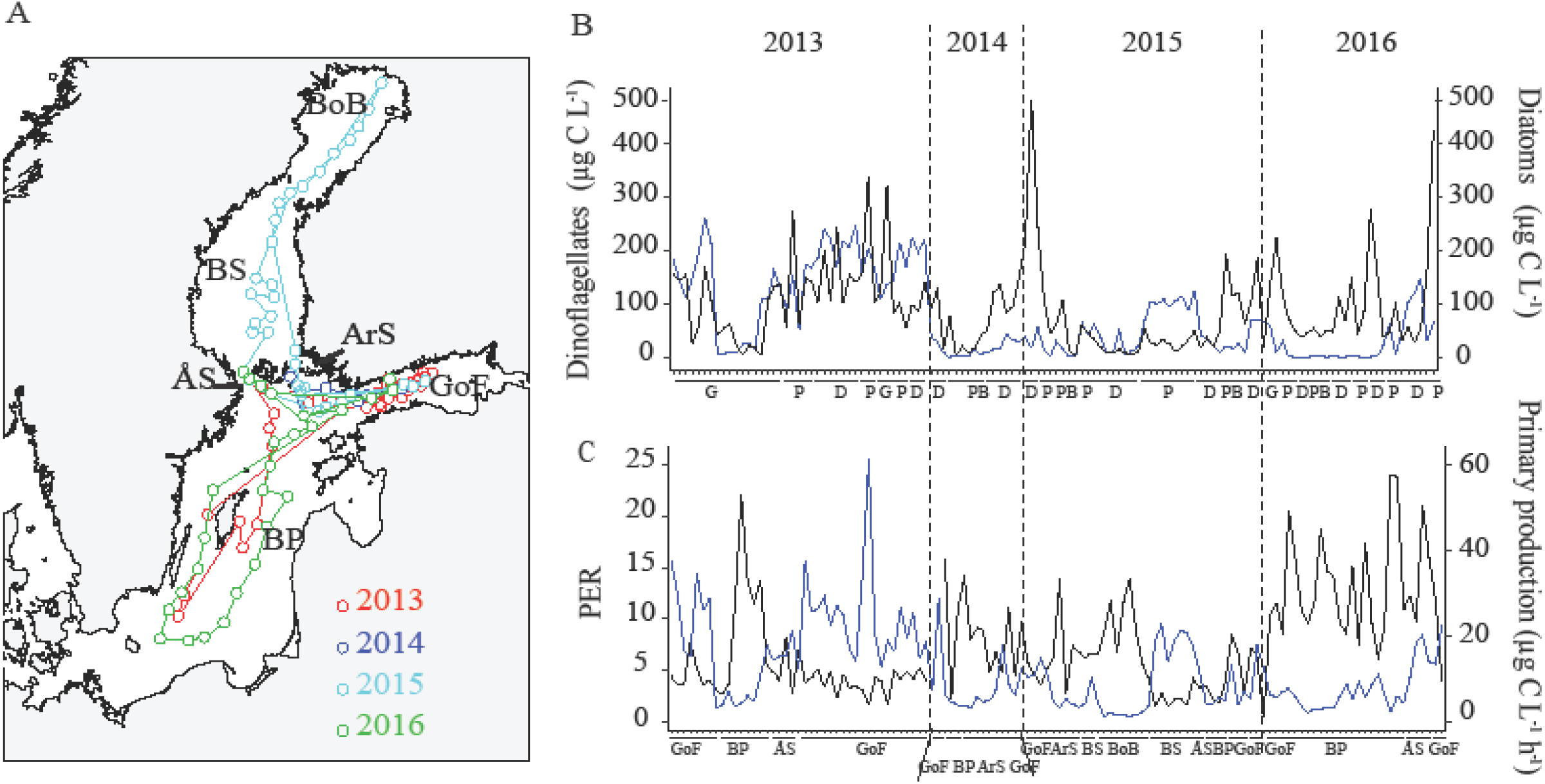
Sampling transects and phytoplankton dynamics across the studied period. A) Map with the sampling transects during the four cruises: 2013 (red), 2014 (dark blue), 2015 (light blue) and 2016 (green). B) Total carbon biomass of dinoflagellates (black) and diatoms (blue) during the cruises and bloom phases. C) Percentage of extracellular release (PER, black) and primary production (PP, blue) in the different sub-basins. See Table 1 for more details of the x-axis abbreviations.

Diatoms and dinoflagellates have different ecological features, as diatoms grow faster than co-occurring dinoflagellates ^11^, they also differ in terms of quality and quantity of dissolved organic matter (DOM) release ^12–14^, and have different sedimentation patterns ^10,15^. Since bacteria are the main consumers of the DOM produced by phytoplankton, also in the Baltic Sea ^1,16^, understanding how the predicted changes in the spring bloom may impact bacterial community composition and activity is essential to anticipate the biogeochemical consequences of this global warming driven process.

Assessing these issues in the Baltic Sea is complex due to the large environmental and spatial-temporal heterogeneity between subbasins, i.e. the pronounced salinity (2 – 20) and temperature gradients (0 – 20°C)^8^. Thus, the taxonomic composition of bacteria varies along the salinity gradient with Alphaproteobacteria and Gammaproteobacteria dominating in more saline waters whereas Actinobacteria, Betaproteobacteria and Planctomycetes are favored by lower salinity ^17–19^ The strong seasonality in the Baltic Sea also shapes bacterial community structure and dynamics: Alphaproteobacteria are present throughout the year ^16,19^, Bacteroidetes, Beta- and Gammaproteobacteria are associated with phytoplankton spring blooms^17,20,21^ and Verrucomicrobia, Actinobacteria and Planctomycetes appear later in the Year^18,22^. Although it is clear that both physical and chemical properties affect bacterial communities, most of these studies have been restricted to specific sub-basins or did not capture the full development of the spring bloom. The potential of diatoms to shape the bacterial communities towards the dominance of Gamma-, Betaproteobacteria and Bacteriodetes and to increase heterotrophic production in the Baltic Sea was recently demonstrated a in mesocosm study^23^. However, so far no study has addressed this issue in natural conditions or across the naturally occurring environmental gradients in the Baltic Sea.

The aim of this study was to investigate how differences in the taxonomic composition of phytoplankton spring bloom communities (diatom- or dinoflagellate-dominated), affect the community structure and dynamics of the associated bacterioplankton across different areas of the Baltic Sea. We collected samples from four consecutive years (2013-2016) during and after the spring bloom, from the northern Gulf of Bothnia to the southern Baltic Proper. Based on our previous observations from laboratory studies^23^, we hypothesized that the predicted increase in dinoflagellates may lead to less productive bacterial communities dominated by less copiotrophic bacterial groups.

## Material and Methods

### Study area, sampling design and environmental variables

The water samples were collected during four cruises (2013-2016) on board of the R/V Aranda. In total, 127 stations were sampled and the area spanned from the southern Baltic Proper (BP, 55°22’N) to the northern Bay of Bothnia (BoB, 65°53’N, Fig 1). At all stations, the water was collected from 3 m depth either using an oceanographic rosette with Niskin bottles of 5 L (n = 122) or from the flow-through system on board (n = 5). Salinity, temperature, and depth, were recorded *in situ*. Inorganic nutrients, measured as nitrogen (NO_2_+NO_3_-N, NH_4_-N), phosphorus (PO_4_-P), and dissolved silica (DSi), particulate organic carbon (POC), biogenic silicate (BSi), dissolved organic carbon (DOC), dissolved organic nitrogen (DON) and chlorophyll *a* (Chl *a)* concentrations were measured as described by Camarena-Gómez, et al.^23^. The particulate organic carbon to chlorophyll *a* (POC:Chl *a)* ratio, based on weight (μg L^-1^), was calculated as an indicator of the trophic status (autotrophic or heterotrophic) of the system, considering heterotrophic status those values larger than 200 ^24^

### Plankton community composition, carbon biomass and productivity

Samples for the determination of nano- and microplankton, including phytoplankton, were determined as described by Lipsewers and Spilling^25^. A modified (carbon biomass instead of biovolume) diatom/dinoflagellate (Dia/Dino) index = diatom biomass/diatom + dinoflagellate biomass, was calculated for each station according to Wasmund, et al.^26^. The values higher than 0.5 indicate diatom dominance whereas values lower than 0.5 indicate dinoflagellate dominance. Primary production (PP) was estimated by measuring the incorporation of ^14^C-labelled sodium bicarbonate (DHI, Denmark) according to Gargas ^27^ and following the method described in Camarena-Gómez, et al.^23^.

The percent extracellular release of ^14^C (PER) was calculated based on the dissolved organic fraction (DO ^14^C < 0.2 μm) of the total ^14^C fixation after 24 h incubation. Since substantial heterotrophic consumption of DOC (on average, 30-50 %) might take place during the incubation period^28^, the results are regarded as net DOC production rates.

Four different stages of the spring bloom were defined based on the concentration of inorganic nutrients (PO_4_^3-^ and NO_3_^-^+NO_2_^-^) and Chl *a* (Table S1): the bloom growth phase (Growth), the peak of the bloom phase (Peak), the declining bloom phase (Decline) and after the bloom phase (Postbloom). The NO_4_^3-^+NO_2_^-^ was used for the N-limited sub-basins Gulf of Finland (GoF), BP, Archipelago Sea (ArS), Åland Sea (ÅS) and Bothnian Sea (BS); and PO4^3-^ for the P-limited BoB subbasin. High Chl *a* was defined as within 20 % of the average peak concentration during spring, which is different in the different sub-basins.

### Bacterial heterotrophic production, abundance and community composition

Bacterial cell production (BPT) and protein production (BPL) were measured in triplicates by using the dual labeling with [metil-^3^H]-thymidine and [^14^C (U)]-leucine (PerkinElmer Inc., Waltham, MA, USA) of samples processed by the centrifugation method as described in Camarena-Gómez, et al.^23^. Bacterial abundances (BA) were determined in duplicates by flow cytometry according to Gasol and Del Giorgio^29^, applying some modifications of the method described in Camarena-Gómez, et al.^23^. Bacterial community analysis including DNA extraction and *Illumina* sequencing were done as described by Camarena-Gómez, et al.^23^. The post-processing of the sequences was done according to the pipeline described by Logares ^30^. The Operational Taxonomic Units (OTUs) were clustered at 99 % of similarity. After the removal of singletons as well as the OTUs classified as chloroplasts and mitochondria, based on the classification with SILVA v123, 2 247 OTUs and 2 250 000 reads were retained. Samples with less than 7 000 reads (4 samples) were removed. The remaining samples (122) were rarefied using *rrarefy* in R to the lowest number of reads (n = 7 024 reads). Downstream analyses were carried out in the rarefied table, including 122 samples and 2 128 OTUs. Raw sequence data will be deposited in the European Nucleotide Archive (acc. nun. XXXXX).

### Statistical analyses

To assess whether there were differences in the diatom and dinoflagellate carbon biomass, between the different phytoplankton bloom phases a non-parametric Wilcoxon rank sum test was applied. This test was also applied to the bacterial diversity in order to test whether there were differences between the two main bacterial community clusters observed. Non-metric multidimensional scaling (NMDS) analysis was done on the Bray-Curtis dissimilarity matrix. Mantel tests with Spearman correlation were applied to study the correlation between the composition of bacterial and phytoplankton communities. Redundancy analysis (RDA) with forward selection was applied to obtain the most important environmental variables that explain bacterial community variance. This analysis included an ANOVA test in each step (permutation test for RDA under reduced model, permutations: free, number of permutations: 9 999) to identify the significant environmental variables *(p* < 0.05). Spearman correlation was used to obtain the correlation between the main bacterial groups and the significant environmental variables as well as the bacterial activity. All figures and analyses were performed with the software RStudio (RStudio Team 2015) with the packages *colorRamps, plotrix, rworldmap, rworldxtra, vegan, ggplot2, ade4, ggpubr* and *Hmisc* and *ComplexHeatmap*.

### Results Inter-annual and spatial variability in environmental variables and bloom dynamics

The environmental conditions varied largely within and between the years due to the location of the different subbasins and time of the sampling (Table 1, Fig. 1A). For instance, salinity ranged from ~ 8 in the southernmost station of the BP to ~2 in the northernmost part of the BoB (Table 1). In general, the NO_2_^-^+NO_3_^-^ was depleted after the bloom with the exception of the P-limited BoB, where the NO_2_^-^+NO_3_^-^ concentration was similar to the stations in the Growth phase and phosphate was low. The lowest dissolved DSi concentration was measured in the GoF in 2014 while the maximum DSi concentration was detected in the Kemi river plume in the northernmost BoB. The DOC and DON were highest in the ArS in 2014, and higher in the GoF and the Gulf of Bothnia compared with the BP.

**Table 1.** Environmental and biological variables measured along the four sampling transects (2013-2016). Values represent the range between the minimum and maximum values, in bold, within the four years. The different sub-basins covered are indicated: Gulf of Finland (GoF), Baltic Proper (BP), Åland Sea (ÅS), Archipelago Sea (ArS), Bothnian Sea (BS), Kvark (Kv) and Bay of Bothnia (BoB). The different phytoplankton bloom phases are indicated: Growth (G), Peak (P), Decline (D) and Post-bloom (PB), defined based on the Chl *a*, nitrate (NO_3_^-^) and phosphate (PO_4_^3-^) concentrations during the sampling at each subbasin (see Methods and Table S1). The inorganic nutrients and dissolved organic nitrogen are presented in μmol L^-1^ and the dissolved organic carbon is presented in mmol L^-1^. The particulate organic carbon to Chl *a* (POC:Chl *a)* ratio indicates the trophic status (considering heterotrophic status those values larger than 200) and the diatoms:dinoflagellates index (Dia/Dino index) indicates diatom dominance or dinoflagellates dominance with values higher or lower than 0.5, respectively.

|  | #of Stations | Subbasins | Bloom phase | Temp (° C) | Salinity (psu) | NO<sub>2</sub><sup>-</sup>-NO<sub>3</sub><sup>-</sup> | NH<sub>4</sub><sup>+</sup> | PO<sub>4</sub><sup>3-</sup> | DSi | DOC | DON | POC:Chl *a* | Dia/Dino index |
| --- | --- | --- | --- | --- | --- | --- | --- | --- | --- | --- | --- | --- | --- |
| **2013**<br>**April**<br>**10th-**<br>**26th** | 29 | GoF | G, P, D | 0.76-1.98 | 5.09-6.38 | 0-5.85 | 0.10-0.39 | 0.19-0.48 | 7.84-19.75 | 0.30-1.08 | 10.06-45.57 | 33.36-52.47 | 0.30-0.86 |
|  | 11 | BP | G, P | 1.62-2.72 | 5.97-7.24 | 0-2.08 | 0.04-0.19 | 0.06-0.33 | 8.27-14.24 | 0.41-1.01 | 13.49-30.78 | 38.96-100.35 | 0.09-0.94 |
|  | 4 | ÅS | G, P | 1.29-1.59 | 5.45-6.16 | 0-2.59 | 0.07-0.13 | 0.02-0.35 | 8.92-13.10 | 0.35-0.51 | 16.07-17.85 | 44.54-56.39 | 0.36-0.74 |
| **2014**<br>**May**<br>**5th-10th** | 9 | GoF | D, PB | 4.25-5.18 | 5.03-6.01 | 0-0.26 | 0.5-0.24 | 0.12-0.33 | 2.27-9.10 | 0.50-1.05 | 16.07-28.70 | 49.43-80.00 | 0.11-0.29 |
|  | 2 | BP | D, PB | 4.62-5.69 | 6.56-6.63 | 0 | 0.05 | 0.28 | 7.60-11.60 | 0.44-0.52 | 15.16-16.06 | 106.82-176.30 | 0.01-0.36 |
|  | 4 | ArS | D, PB | 4.58-5.17 | 5.90-6.29 | 0-0.4 | 0.08-0.11 | 0.20-0.33 | 6.01-8.10 | 0.60-1.12 | 16.02-67.86 | 88.28-208.84 | 0.05-0.49 |
| **2015**<br>**May**<br>**4th-15th** | 5 | GoF | D, PB | 3.91-5.35 | 4.96-5.86 | 0-0.02 | 0.08-0.44 | 0.18-0.36 | 5.50-11.20 | 0.46-1.07 | 16.36-25.40 | 75.28-119.28 | 0.12-0.40 |
|  | 6 | ArS | P, PB | 4.41-6.09 | 5.75-6.38 | 0-0.23 | 0.06-0.13 | 0.06-0.33 | 5.60-11.10 | 0.32-0.64 | 11.33-18.58 | 58.81-190.20 | 0.04-0.32 |
|  | 4 | ÅS | D | 5.03-6.22 | 5.41-6.06 | 0 | 0-0.06 | 0.05-0.22 | 6.70-8.70 | 0.37-0.47 | 12.73-13.97 | 76.75-108.31 | 0.17-0.68 |
|  | 4 | BP | D, PB | 4.52-5.69 | 5.72-6.56 | 0-0.44 | 0.05-0.34 | 0.27-0.40 | 7.6-11.8 | 0.37-0.55 | 15.16-18.95 | 83.49-149.96 | 0.09-0.33 |
|  | 10 | BS | P | 3.59-4.16 | 5.12-5.54 | 0-0.38 | 0.05-0.13 | 0-0.21 | 8.10-12.70 | 0.33-1.07 | 12.40-18.66 | 31.93-82.12 | 0.48-0.90 |
|  | 9 | Kv, BoB | P, D | 1.35-4.73 | 2.20-5.09 | 0-7.47 | 0.04-0.35 | 0-0.02 | 9.40-55.70 | 0.35-0.68 | 15.73-55.53 | 70.71-223.37 | 0.36-0.68 |
| **2016**<br>**April**<br>**4th-15<sup>th</sup>** | 4 | GoF | G, P, D | 1.56-2.89 | 4.95-5.20 | 0-6.64 | 0.14-0.22 | 0.34-0.61 | 12.21-19.09 | 0.44-0.58 | 14.71-31.52 | 25.85-57.10 | 0.13-0.77 |
|  | 20 | BP | G, P, D, PB | 2.47-5.90 | 5.92-7.77 | 0-1.89 | 0-1.19 | 0.16-0.57 | 11.77-17.65 | 0.32-0.62 | 10.73-22.10 | 44.49-260.31 | 0.01-0.58 |
|  | 4 | ÅS | P | 2.84-3.28 | 5.51-5.68 | 0-0.17 | 0.11-0.18 | 0.12-0.18 | 15.57-18.17 | 0.34-0.99 | 12.40-14.54 | 58.41-67.04 | 0.61-0.80 |

Overall, diatom carbon biomass was significantly higher in the Growth and Peak phytoplankton bloom phases compared to the Decline and Postbloom phases (Wilcoxon test, *p* < 0.05), whereas dinoflagellate carbon biomass was also significantly high in the Decline phase (Wilcoxon test, *p* < 0.05). However, no clear increasing trend in the biomass of dinoflagellates compared to diatoms was observed across the studied years. In 2013, we sampled right after the ice melt in the Gulf of Finland and the water temperature was low in the entire sampling area (Table 1). The spring bloom was in the Growth to Decline phase in the 2013 cruise (Table 1), and was largely dominated by diatoms, displaying much higher carbon contribution to total biomass than in the other three cruises (Fig. 1B). In 2014, the temperature was 4-5.5 °C and the spring bloom was in a more advanced phase with low phytoplankton biomass (Fig. 1B). In 2015, the conditions were similar to 2014, with the exception of the Gulf of Bothnia where the spring bloom was in the Peak to Decline phase at the time of the sampling. In 2016, we sampled the same areas and time of the year as 2013, but the temperature was warmer in the BP subbassin and the spring bloom was more advanced in the latest year than in the first sampling cruise (Table 1). In these warmer cruises, dinoflagellates contributed > 300 μg C L^-1^ to the carbon biomass at some stations (Fig. 1B).

High PP was observed during the Growth and Peak phases (Fig. 1C), following the tendency of the Chl *a* and coinciding with the stations that presented high carbon biomass (Fig. 2A, Table S2). The PER, measured after 24 h, was high in the Decline and Postbloom phase in the BP and the BoB, in contrast to PP (Fig. 1C, Table S2). The trophic status, estimated as the POC:Chl *a* ratio, was in general low, except at some stations in the BP and the BoB (Table 1).

**Figure 2.**
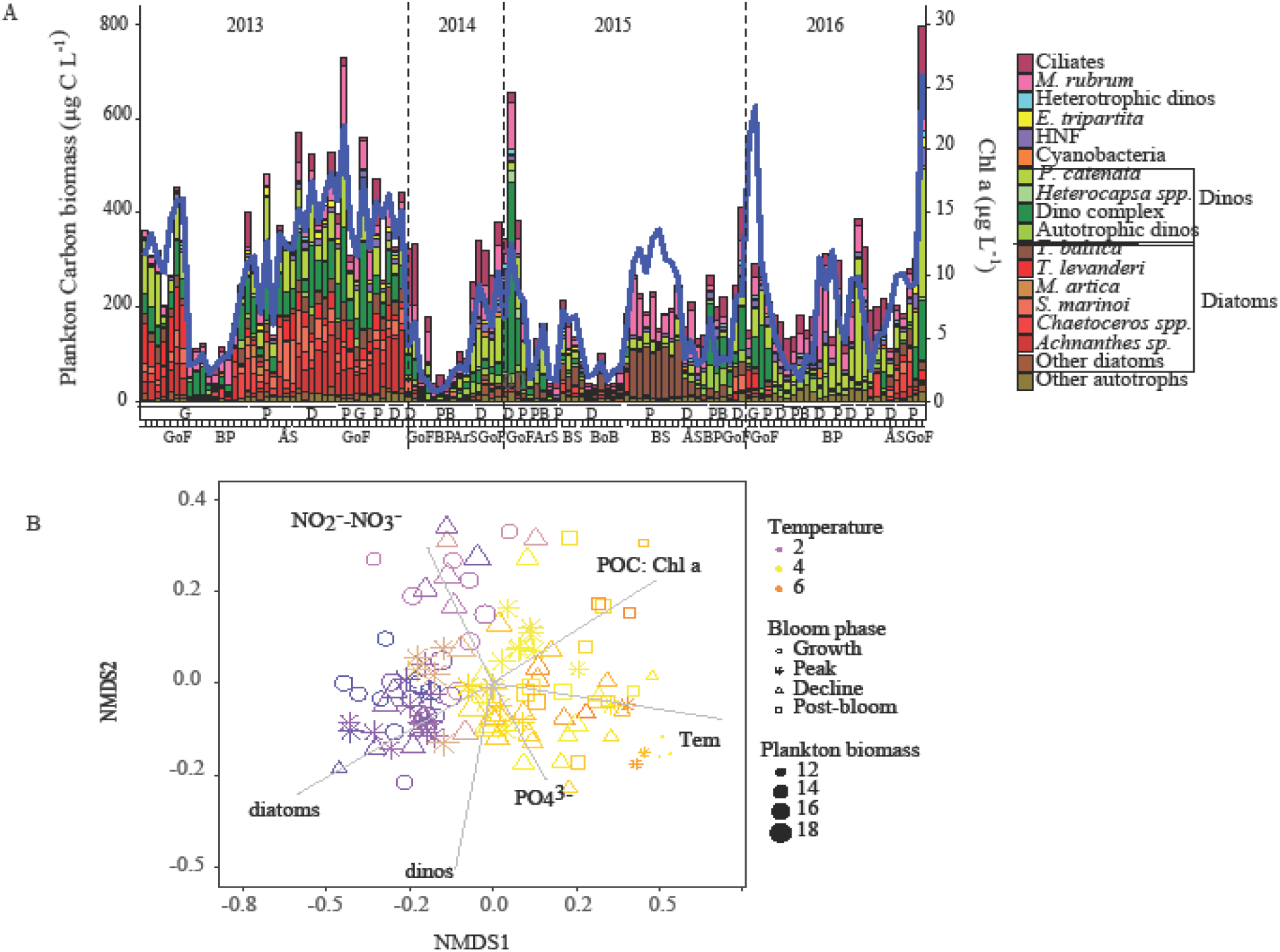
Plankton dynamics in A) total carbon biomass of the main plankton groups and the chlorophyll *a* (Chl *a)* concentration at each station. The years are indicated on top of the figure and the bloom phases and the sampled subbasins are at the bottom of the figure. See Table 1 for more details of x-axis abbreviations. B) Non-metric multidimensional scaling (NMDS) plot based on Bray-Curtis distances between plankton community taxonomic composition (N = 126, stress = 0.1602). The vectors indicate the environmental factors most strongly explaining the observed distribution based on RDA forward selection (NO _2_^-^-NO_3_^-1^; nitrate-nitrite, Tem; temperature, PO_4_^3-^; phosphate, POC:Chl *a;* particulate organic carbon to Chl *a* ratio, diatoms and dinos; total carbon of diatoms and dinoflagellates). The colour gradient indicates the water temperature, shape indicates the different bloom phases and size indicates the log-transformed total carbon biomass.

### Changes in plankton community composition

We observed pronounced differences in taxonomic composition of the studied plankton communities (mainly phytoplankton and nano-microzooplankton here) across the different subbasins and years, which appeared mainly shaped by the inorganic nutrient concentrations and the water temperature (Fig. 2A, B). Diatoms largely dominated in the GoF and ÅS (Table 1), with high contributions of *Achnanthes taeniata, Skeletonema marinoi, Thalassiosira levanderi*, and *Chaetoceros* spp., whereas in the Bothnian Sea, *T. baltica* dominated during the Peak bloom phase (Fig. 2A). Dinoflagellates were dominated by the dinoflagellate complex, mainly formed by *Biecheleria baltica* and/or *Gymnodinium corollarium* in the GoF and BP. The chain forming *Peridiniella catenata* was present in all the subbasins except at the stations with clearly low carbon biomass (Fig. 2A). Other autotrophic dinoflagellates *(Heterocapsa* spp.) and heterotrophic/mixotrophic dinoflagellates (e.g. *Protoperidinium* spp. and *Dinophysis* spp., respectively) were also present, but their contribution to the total plankton carbon biomass was < 5 %. The contribution of other autotrophic plankton organisms, such as Crytophyceae, Chrysophyceae, Prasinophyceae, Prymnesiophyceae, Chlorophyceae, to the total carbon biomass was also generally low (< 2 μg C L^-1^).

The mixotrophic ciliate, *Mesodinium rubrum*, was present across all sub-basins having highest contribution in the BS and being in general more abundant in 2016 compared with other years (Fig. 2A). Heterotrophic ciliates were most abundant during the late phase of the bloom in the GoF and the northern BP. The silicoflagellate *Ebria tripartita* and the heterotrophic nanoflagellates (HNF) had generally low abundance but contributed more than 10 μg C L^-1^ to the total carbon biomass during the Peak and Decline phases of the bloom also at some stations in the GoF and the BP.

### Bacterial community composition and its environmental and biological drivers

The taxonomic composition of the studied bacterial communities varied largely across the stations sampled (Fig. 3). Overall, Alphaproteobacteria, mainly SAR11, dominated in all the sampled years (Fig. 3A). Bacteroidetes were dominated by the class Flavobacteriia, which comprised on average ~ 25 % of the total relative abundance. Within Flavobacteriia, the genus *Flavobacterium* contributed 8-12 % to the total relative abundance during the Peak/Decline phase in 2013 and 2016, together with *Fluviicola, Polaribacter* and NS3 marine group, whereas *Owenweeksia* was found in the Growth phase. Gammaproteobacteria *(Crenothrix)* had higher relative abundance (> 10 %) in the GoF and the ÅS in 2013 and 2016. The classes Actinobacteria (hgcl clade) and Acidimicrobiia (CL500-29 marine group) had higher relative abundance in 2014 and 2015. The phylum Phycisphaerae (CL500-3) was scarce but reached its highest values (~ 8 %) in the Gulf of Bothnia in 2015.

**Figure 3.**
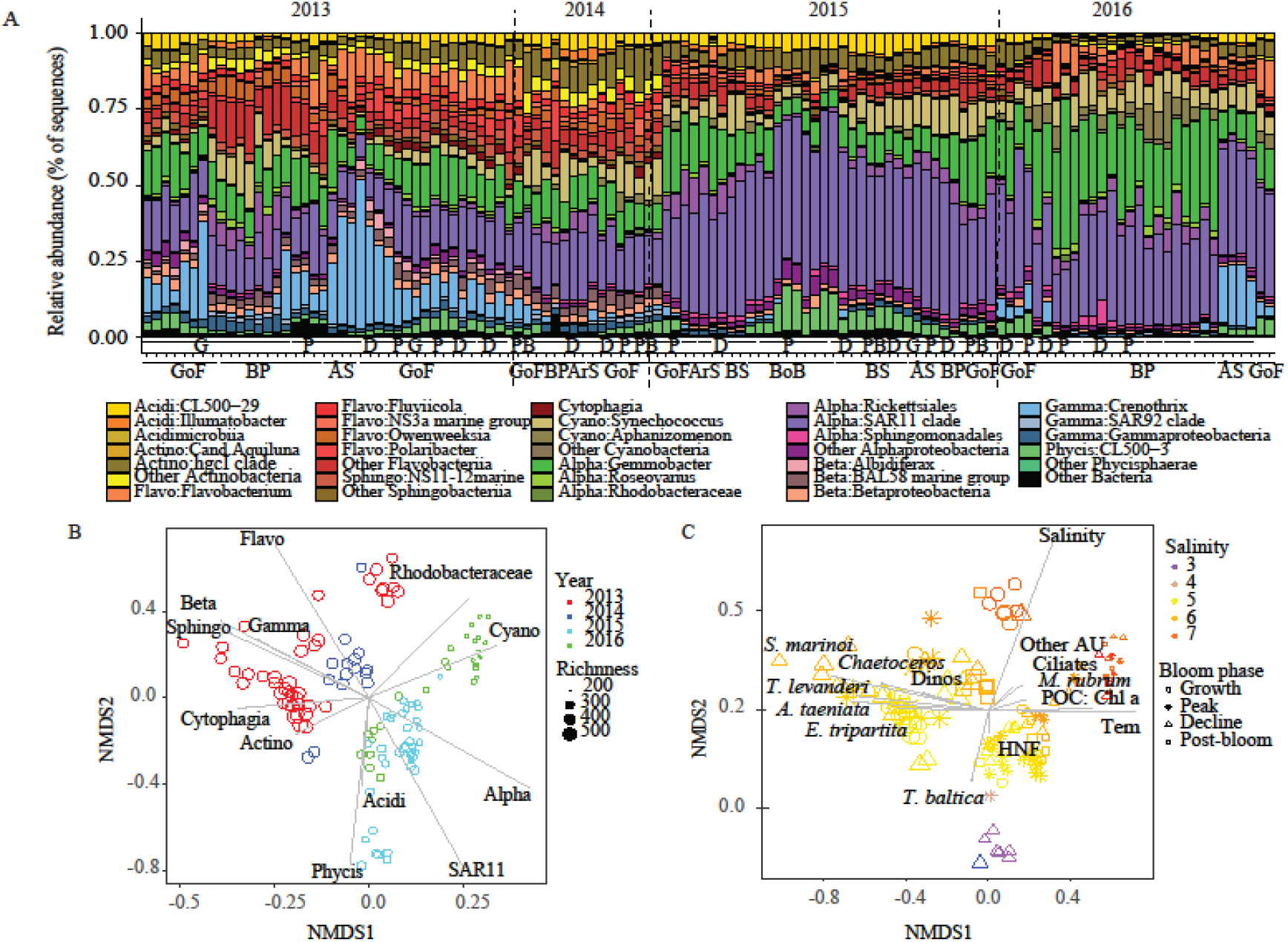
Bacterial community dynamics and their environmental drivers. A) Relative abundance of the bacterial groups (class-family and genus level) that contribute more than 0.5 % of the total OTUs. Abbreviations as in figure 1. B) Non-metric multidimensional scaling (NMDS) plot based on Bray-Curtis distances of bacterial community structure (N = 122, stress = 0.134). Colour-code indicates the years (see Fig. 1) and size indicates the OTU richness in each community. Vectors indicate the main bacterial groups found in the four years. The abbreviations in the legend of Fig. A) and in Fig. B) indicate the main clades found during the sampling: Acidi; Acidimicrobiia, Actino; Actinobacteria, Flavo; Flavobacteriia; Sphingo; Sphingobacteriia; Cyano; Cyanobacterias, Alpha; Alphaproteobacteria; Beta; Betaproteobacteria; Gamma; Gammaproteobacteria; Phycis; Phycisphaerae (Phy). C) Non-metric multidimensional scaling (NMDS) plot linking bacterial community structure with environmental variables and plankton community. Colour-code indicates the salinity gradient and shape indicates the phytoplankton bloom phase. The vectors indicate the significant environmental variables as follow: *Achnanthes taeniata, Skeletonema marinoi, Thalassiosira levanderi, Chaetoceros* spp., Dinos (dinoflagellate complex), *Ebria tripartite, T. baltica*, HNF (heterotrophic nanoflagellates), temperature, POC Chl *a, Mesodinium rubrum*, ciliates, other autotrophs and salinity.

Based on their overall taxonomic structure, the communities were clustered into two main groups (Wilcoxon; *p* < 0.001) differing largely in taxonomic richness (Fig. 3B) and diversity (Shannon index; table S2) as well as in the bacteria abundances of the major bacterial groups. The group with high taxonomic diversity comprised communities sampled during 2013 and 2014, and showed higher relative abundances of Bacteroidetes, Gammaproteobacteria and Actinobacteria, whereas the low taxonomic richness group (2015-2016) showed a dominance of Alphaproteobacteria (SAR11 and Rhodobacteraceae) and Cyanobacteria (Fig. 3B).

Although the bacterial community structure was strongly influenced by the salinity gradient (Fig. 3C), the shift in community structure observed between 2013-2014 and 2015-2016 seemed to be largely driven by certain phytoplankton groups such as the diatoms *A. taeniata, S. marinoi, T. levanderi* and *Chaetoceros* spp., as well as temperature and the POC:Chl *a* ratio (Fig. 3C). The role of the phytoplankton community composition in shaping bacterial assemblages was further supported by the strong correlations between bacterial and phytoplankton taxonomic dissimilarity matrices found across the four years of study (Spearman r values 0.6-0.8, Mantel test *p* = 0.0001; Table S3).

When exploring the individual correlations between different environmental and biological factors and the abundances of the main bacterial groups (Fig. 4), we observed two groups of bacterial taxa showing fundamentally contrasting relationships. Bacteroidetes (Flavobacteriia, Cytophagia and Sphingobacteriia), Gamma- and Betaproteobacteria were strongly correlated with diatoms and *E. tripartita*, and negatively correlated with temperature and POC:Chl *a* ratios (Fig. 4). In contrast, Rhodobacteraceae, SAR11 and Cyanobacteria correlated positively with temperature, POC:Chl *a* ratios, HNFs and ciliates, whereas the dominance of certain diatoms or dinoflagellates caused decreases in their abundances (Fig. 4). Phycisphaerae correlated strongly with the diatom *T. baltica*. Within dinoflagellates, the dinoflagellate complex was the only group that correlated positively with all the bacterial taxa dominant in 2013-2014, but also with Rhodobacteraceae (Fig. 4), which was more prevalent in 2015-2016 (Fig. 3B).

**Figure 4.**
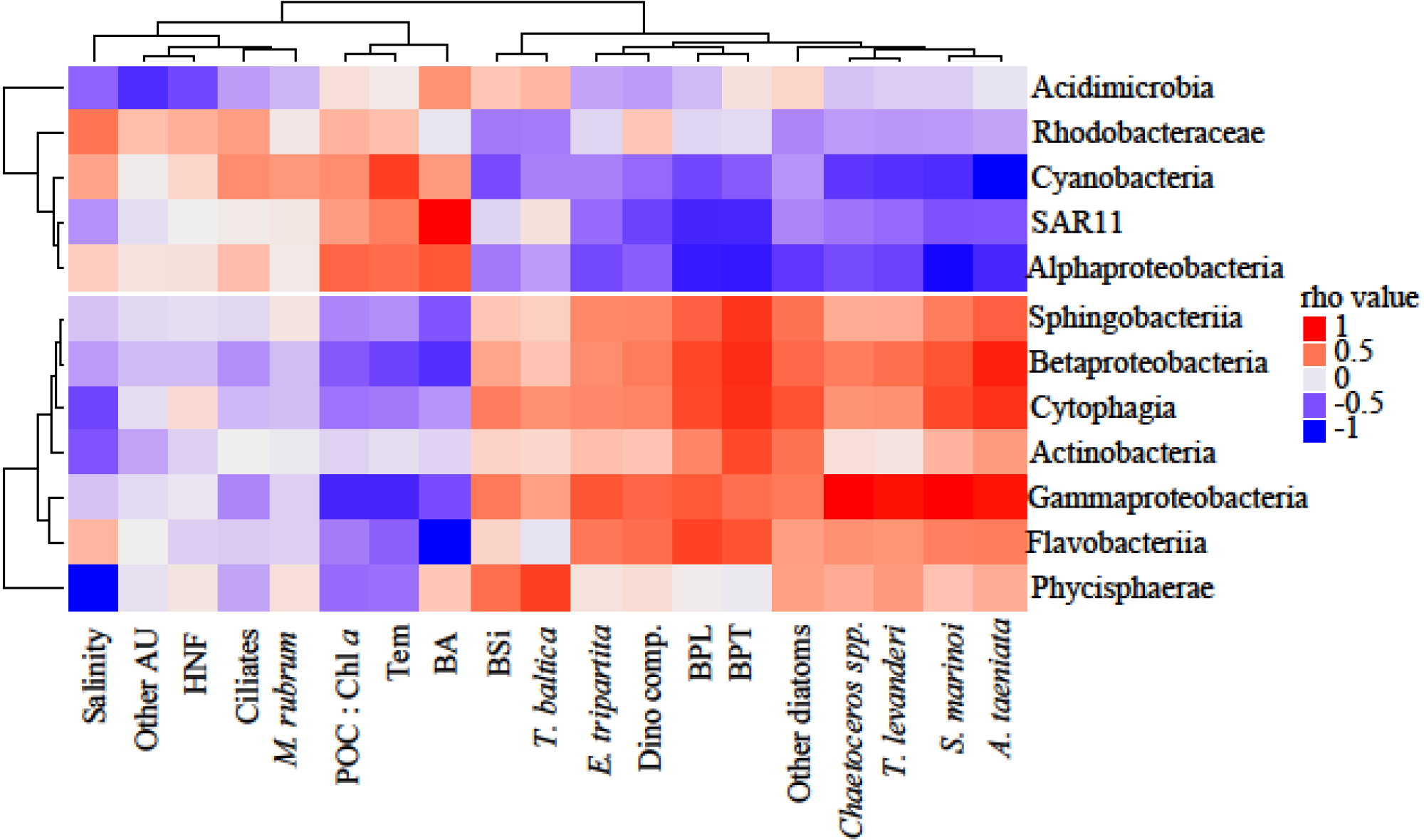
Heatmap with Spearman correlations between significant environmental variables, phytoplankton or heterotrophic plankton groups and the main bacterial groups. Colour-code indicates the correlation coefficients (rho values). The environmental factors (columns) are as in Fig. 3. In addition, BA = bacterial abundance; BSi = biogenic silicate; BPT = bacterial production thymidine and BPL = bacterial production leucine. N = 122.

The observed changes in composition of the phytoplankton communities were accompanied by a large spatial-temporal variation of bacterial abundances and production (Fig. 5). Overall, the bacterial production rates measured with leucine (BPL) were highest in 2013 (Fig. 5A, Table S2), having similar pattern of the PP (Fig. 2A), while when measured with thymidine (BPT), the rates were highest in 2014 (Fig. 5A). In contrast, the highest BA values were recorded at the stations sampled during the 2015 cruise (Fig. 5B; Table S2).

**Figure 5.**
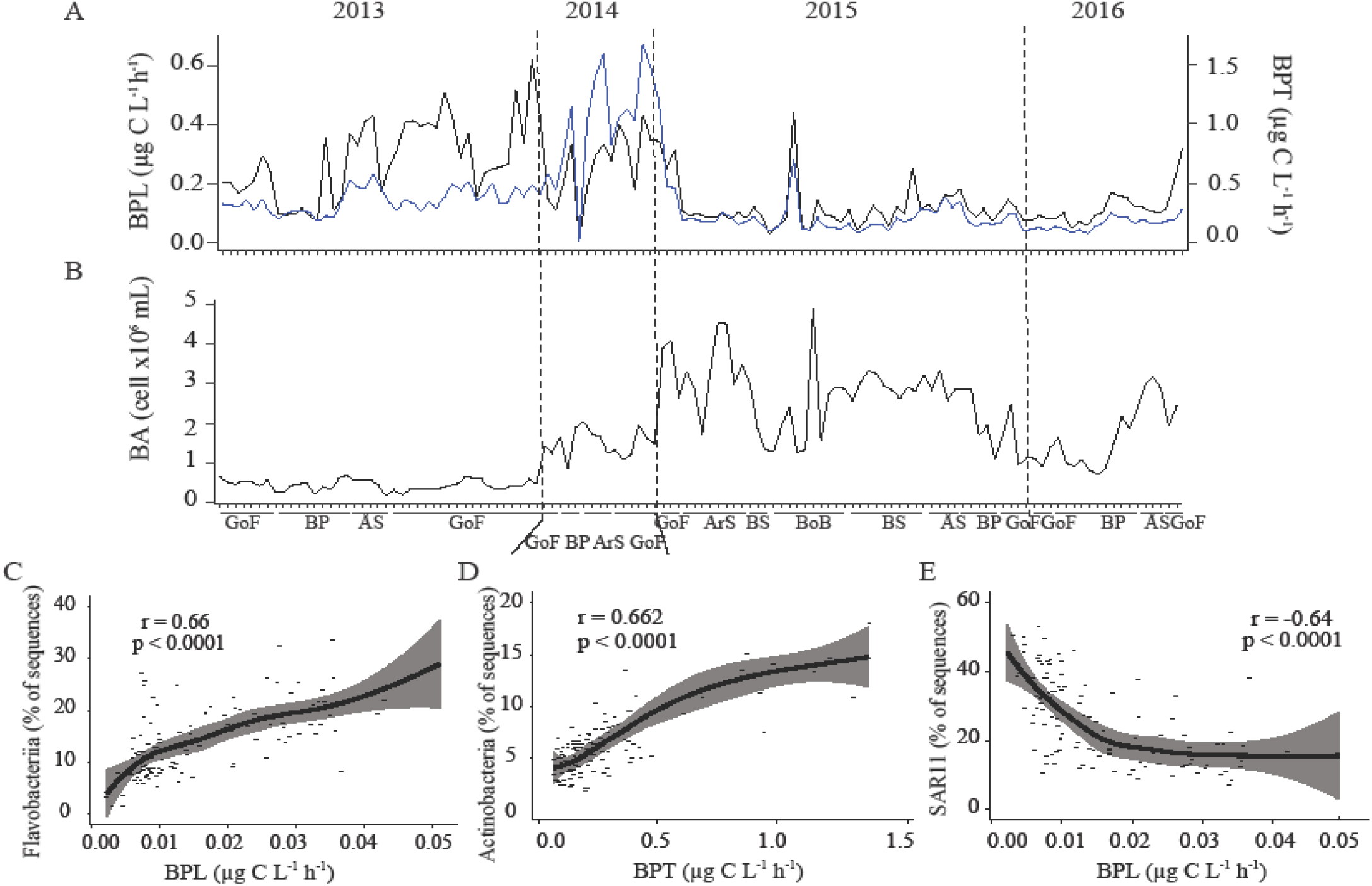
Bacterioplankton productivity and abundance during the sampled spring blooms and the link to bacterial community. A) Bacterial heterotrophic production based on leucine incorporation (BPL, black) and thymidine incorporation (BPT, blue), and B) bacterial abundance. The x-axis indicates the sub-basins across the sampling. See Fig. 1 for more detail of legend abbreviation. The dashed lines separate the different years. C-E) Spearman correlations between bacterial production measured as leucine uptake (BPL) or thymidine uptake (BPT) and the relative abundance (%) of Flavobacteriia (C), Actinobacteria (D) and SAR11 (E), during the study. Rho values and p values are presented on the figure. N = 122; p < 0.05.

The differences in BCC had an impact on community functioning. Most of the bacterial groups that were prevalent in the high diversity assemblages from 2013 and 2014, i.e. Beta-, Gammaproteobacteria, Actinobacteria, Bacteroidetes, and more specifically Flavobacteriia showed strong positive correlations with BPL and BPT (Fig. 5C-D; Table S4). In contrast, the abundance of Alphaproteobacteria (SAR11 in particular), was negatively correlated with bacterial production rates (Fig. 5E, Table S4).

## Discussion

Our results demonstrate that the bacterioplankton community structure was strongly affected by the different phytoplankton communities, either dominated by diatoms or dinoflagellates, and by the different phases of the phytoplankton bloom. Although our results do not evidence a clear trend of increasing dominance of dinoflagellates from 2013 to 2016, we did capture much higher carbon biomass of diatoms during the 2013 cruise than in the other years. This heterogeneity allowed us to explore how changes in the dominant phytoplankton groups are translated into changes in the structure and functioning of the associated bacterioplankton communities.

### Dynamics of environmental variables and phytoplankton biomass

We found that the different environmental conditions, primarily temperature and inorganic nutrient concentrations such as NO3^-^ and PO_4_^3-^, shaped the phytoplankton community structure and bloom dynamics in terms of Chl *a* concentration, PP and dominant species, in agreement with previous studies in the Baltic Sea^9,31^. The phytoplankton community during the early bloom development, with low temperature and high inorganic nutrient concentrations, was clearly dominated by diatoms, in particular in the GoF. Diatom dominance was also associated with high primary productivity suggesting an actively growing community. The prevailing diatoms were the taxa that typically occur during the cold water season ^32–34^. In the BS, the Peak phase was dominated by the single diatom species *Thalassiosira baltica* along with high carbon biomass of the mixotrophic ciliate *Mesodinium rubrum*, which both seem to be common in this sub-basin ^7^. The diatom-dominated communities co-occurred to a varying extent with several dinoflagellates, most likely *Gymnodinium corollarium* and *Biecheleria baltica* ^35,36^ in the GoF and the BP, similarly to the earlier observations^2,3^. The heterotrophic *Ebria tripartita* was more abundant during diatom dominance, likely because it grazed on diatom species such as *Skeletonema* sp. and *Thalassiosira* spp. ^37^.

During the declining phytoplankton bloom, the water was warmer, the Chl *a* concentration and the total carbon biomass were lower and dinoflagellates contributed more to the biomass than in the Growth and Peak phases. This is the typical situation in the GoF with diatoms being more abundant in early spring and dinoflagellates gradually becoming more dominant as the nutrients are depleted^10,38^. For instance, the stations sampled during 2014 presented low phytoplankton biomass and also low DSi concentration in the GoF, suggesting that a diatom-dominated bloom had settled out of the surface layer before the time of sampling. This probably explained the similarity in structure between the bacterial communities and productivity found in 2014 and those in the diatom-dominated stations of 2013. In the BoB, the low phytoplankton carbon biomass was limited by phosphorus with excess NO3^-^ and DSi and high DOC and DON concentrations. This sub-basin is P-limited ^9^ and is more turbid due to terrestrial DOM inputs, which limits light penetration resulting in light-limited primary production ^39–41^.

The abundance of nano- and microzooplankton (HNF, ciliates and *M. rubrum)* were also higher during the Decline phase of the bloom, which, together with higher POC:Chl *a* ratios (> 175), points to a more heterotrophic plankton community^24^. A decrease in phytoplankton biomass and an increase in the proportion of dinoflagellates and heterotrophic organisms have been projected for a future, warmer Baltic Sea ^1^, in addition to the earlier onset of the spring bloom^42^. Our results represent different temperature scenarios, for example the 2013 and 2016 cruises sampled the same subbasins during varying environmental conditions with 2016 being warmer and the bloom in a more advance stage, in line with these predictions.

### Bacterial community dynamics and their environmental drivers

The changes in the phytoplankton community composition between the bloom phases resulted in pronounced shifts in the structure of the associated bacterial communities and likely related to the availability of the released DOM, largely produced by phytoplankton ^43,44^ The availability of this DOM can lead to changes in bacterial community dynamics ^45–47^, which have been also reported in several field studies during phytoplankton spring blooms ^16,17,48^

The studied bacterial communities clustered into two clearly different groups of high and low taxonomic richness and diversity, which corresponded to the communities sampled during 2013-2014, and those sampled during 2015-2016, respectively. Salinity and temperature appeared as relevant factors in shaping the bacterial communities in the Baltic Sea, similar to previous studies ^20,22,49^. However, in our study, the salinity was not responsible for the segregation observed in the bacterial communities. Instead, changes in the carbon biomass of specific phytoplankton groups such as *Achnanthes taeniata, Chaetoceros* spp., *Skeletonema marinoi*, and *T. levanderi* emerged as some of the most important drivers in the configuration of bacterial communities. Thus, the diatom bloom observed in 2013 and the diatom bloom that had likely recently settled out of the surface layer in 2014 promoted a higher diversity of associated bacterial communities with more Gammaproteobacteria, Betaproteobacteria, and Bacteroidetes. This conclusion was further strengthened by the strong and positive correlations between the diatom species and these copiotrophic bacteria, in accordance with their preference for productive or bloom-like conditions. All these groups are known to be favored during diatom blooms ^23,50^. In contrast, the lower diatom carbon biomass observed during 2015 and 2016 were associated with less diverse communities dominated by the SAR11 group in 2015 and also by Rhodobacteraceae (genus *Gemmobacter)* in 2016, supported by the strong negative correlations between this bacterial community structure and the phytoplankton carbon biomass. The conditions within the less productive stations sampled during 2015 and 2016 likely promoted the dominance of SAR11, which typically occurs in nutrient-poor environments^51,52^. Other groups like Phycisphaerae, and in particular CL500-3, appeared to follow more clearly the salinity gradient and was strongly associated to the presence of *T. baltica*. The presence of Phycisphaerae has been associated to the lower salinity of the Gulf of Bothnia and to a higher influence of allochthonous DOM ^19^ Interestingly, temperature emerged as a one of the main drivers for the structure of both, the phytoplankton and bacterioplankton communities, suggesting that the predicted increases in water temperature along the Baltic Sea ^53,54^ may lead to dramatic changes in the planktonic assemblages inhabiting this vulnerable system.

Within the high-diversity bacterial communities found in 2013 and 2014, we observed that the dynamics of Gammaproteobacteria and Bacteroidetes differed between bloom phases and subbasins. Bacteroidetes were always present with fluctuating relative abundance, whereas Gammaproteobacteria, dominated by the genus *Crenothrix*, peaked shortly and were restricted to those stations with complex diatom communities. These bacterial groups are known to occupy different niches in terms of metabolic affinities: Flavobacteriia are capable of degrading high molecular weight compounds, such as proteins, chitin and polysaccharides ^48,55^, whereas Gammaproteobacteria are more specialized on breaking down low molecular weight substrates, peaking shortly during increased organic and inorganic substrate availability ^56,57^. Thus, their different patterns were likely related to the DOM availability and niche partitioning during the highly dynamic phytoplankton blooms. This was supported by the lower net PER during diatom dominance compared with dinoflagellate dominance, suggesting that diatoms release more labile DOC, and therefore more rapidly consumed, than dinoflagellates (Lignell 1990).

The succession of taxa within Bacteroidetes in the different phases of the bloom was similar to that observed in a mesocosm experiment simulating diatom and dinoflagellate blooms with Baltic Sea bacterial communities^23^. In the Growth phase, *Owenweeksia, Flabovacterium* and *Fluviicola* were more dominant, whereas the genera *Polaribacter* and NS3 marine group were favored in the declining phase. Within Gammaproteobacteria, the presence of *Crenothrix* in surface waters was surprising since this taxon is a methane-oxidizer specialized in the uptake of single-carbon substrate, such as methane and methanol, but also acetate and glucose ^58^. This filamentous bacterium may have benefited by the increase of the suitable substrates, either by fresh and labile DOM released or as a product from a metabolic cascade after the degradation process carried out by other bacterial groups.

A lower diversity of Bacteroidetes was observed in the presence of the diatom *T. baltica* in the BS. A previous study has reported the production of certain polyunsaturated aldehydes (PUAs) by *Thalassiosira spp*. ^59^, which can affect the bacterial protein production of Bacteroidetes negatively ^60^. This could have played a role in the reduced growth of this bacterial group. Any anti-bacterial effects of phytoplankton are not well studied, but there are indications that some species inhibit bacterial growth ^50^.

### Links between bacterial community composition and ecosystem functioning

The observed changes in bacterial community structure resulted in pronounced changes in the bacterial production of the studied communities. Most stations sampled during 2013 and 2014 showed much higher rates of BPL compared to the stations sampled in 2015 and 2016 and were similar or, in some cases, larger than those reported by previous studies conducted at similar temperatures during the spring in the Baltic Sea ^16,61^. The larger values of BPL coincided with the higher abundances of copiotrophic groups such as Betaproteobacteria, Gammaproteobacteria and Bacteroidetes associated to diatom dominance. The variation in BPL between studies conducted at similar temperatures may be due to the different dominating phytoplankton groups that produce DOM of different lability. This emphasizes the importance of the phytoplankton taxonomical composition as the driver of the bacterial carbon cycling mediated by the presence of specific bacterial taxa. Large peaks in BPT were also observed in 2014 indicating an actively dividing community and mostly related to the increases in relative abundance of Actinobacteria, which is known to occur after phytoplankton blooms ^20^ and correlate positively with thymidine incorporation ^62^ In contrast, SAR11 correlated negatively with BPL, which agrees with the small and slow-growing pattern of this clade typically occurring in oligotrophic ecosystems^51,52^. Similar bacterial responses were observed in a mesocosm study, where the growth of diatoms resulted in higher bacterial production and higher dominance of Flavobacteriia, Gammaproteobacteria and Cytophagia than in the presence of dinoflagellates^23^. The robustness of the observed patterns in our study, linking BCC and functioning, are essential in microbial ecology since the studies that explore these relationship are scarce in the Baltic Sea ^63^.

Gammaproteobacteria and Bacteroidetes are known to be quickly grazed by HNF ^51^, which prefer to graze on actively growing bacteria. This may explain the low bacterial abundance of the active community observed in 2013. The dinoflagellate complex also correlated positively with the relative abundances of these bacterial groups. However, previous experimental observations reported a decrease in bacterial productivity upon the addition of *Biecheleria baltica* ^23^, one of the species forming the dinoflagellate complex.

### Conclusions

Our results during and after phytoplankton spring blooms highlight the differences in BCC, diversity and productivity under distinct phytoplankton communities dominated by diatoms and/or dinoflagellates. Diatoms like *Skeletonema marinoi, Achnanthes taeniata* and *Chaetoceros* spp. likely release high-quality DOM that boosts the bacterial production and enhances the development of highly diverse bacterial communities dominated by Gamma-, Betaproteobacteria and Bacteroidetes. In contrast, less diverse and less productive bacterioplankton communities dominated by SAR11 and Cyanobacteria are observed in presence of more dinoflagellates. Based on our results, it is not the overall dominance of dinoflagellates or diatoms what drives the largest changes in bacterial community structure, but rather the presence of particular taxonomic groups. Finally, although temperature appears as a main factor shaping both bacterial and phytoplankton communities, higher temperatures are not necessarily related to higher bacterial production rates. Thus, it is needed to incorporate knowledge on the changes in plankton community composition to be able to understand and predict the biogeochemical consequences of expected temperature increases and associated changes in phytoplankton blooms in Baltic Sea waters.

## Acknowledgements

This study was funded by the Walter and Andrée de Nottbeck Foundation and the Academy of Finland (decision numbers 259164 and 292711). We would like to thank the staff at Tvärminne Zoological Station and the staff on board the Aranda vessel for help with the analytical measurements and sampling during the experiments.

