## Supplementary tables for "Bacterioplankton dynamics driven by inter-annual variation in phytoplankton spring bloom communities in the Baltic Sea" (from another version of this preprint: version 2, 2019-01-09)

**Supplementary figures**

**Table S1.** Definition of the phytoplankton bloom phase based on the concentrations of Chl *a* and inorganic nutrients (PO_4_^3-^ and NO_3_^-^, see Methods). The bloom phases are: growth (Growth), peak (Peak), decline (Decline) and post-bloom (Post-boom) phases. Sub-basins are Gulf of Finland (GF), Baltic Proper (BP), Åland Sea (ÅS), Archipelago Sea (ArS), Bothnian Sea (BS) and Bay of Bothnia (BoB). The different levels of Chl *a* concentrations (µg L^-1^) for each sub-basin are set on the table. The PO_4_^3-^ was in the Bay of Bothnia and the NO_3_^-^ in the rest of the Baltic Sea.

| **Stage spring bloom** | **Growth** | **Peak** | **Decline** | **Post-bloom** |
| --- | --- | --- | --- | --- |
| **Chl *a* concentration** | low-medium | High | medium | low |
| **Sub basin**  GF | <16 | >16 | 4-16 | <4 |
| BP | <8 | >8 | 3-8 | <3 |
| ÅS, ArS, BS | <6 | >6 | 2-6 | <2 |
| BB | <5 | >5 | 1-5 | <1 |
| **Inorganic nutrient concentration** | medium-high | low-medium | low | low |
| **Sub-basin**  GF, BP, ÅS, ArS, BS (NO_3_) | >0.5 | <0.5 | <0.2 | <0.2 |
| BoB (PO_4_) | >0.3 | <0.3 | <0.1 | <0.1 |

**Table S2.** Biological variables measured in the four sampling transects (2013−2016). Values represent the range between the minimum and maximum, indicated in bold, in the sub-basins sampled in each year The variables are chlorophyll *a* concentration (Chl *a*), primary production (PP), percentage of net extracellular release (PER) during 24 hours incubation, bacterial production based in thymidine (BPT) and leucine (BPL), bacterial abundance and the Shannon index of the bacterial community (n = 122).

|  | Subbasins | Chl *a*  µg L^-1^ | PP  µg L^-1^ h^-1^ | PER | BPT  µg L^-1^ h^-1^ | BPL  µg L^-1^ h^-1^ | BA cell x10^6^ mL | Shannon index |
| --- | --- | --- | --- | --- | --- | --- | --- | --- |
| 2013 | GoF | 10.26-21.96 | 13.01-**61.18** | 0.58-3.62 | 0.26-0.51 | 0.16-**0.62** | **0.20**-0.67 | 2.91-**4.61** |
| April | BP | 2.22-11.54 | 3.16-17.64 | 1.92-6.62 | 0.18-0.52 | 0.07-0.33 | 0.23-0.69 | 3.73-4.38 |
| 10^th^-26^th^ | ÅS | 6.10-12.73 | 12.28-21.32 | 1.39-2.08 | 0.43-0.57 | 0.17-0.43 | 0.46-0.59 | 3.83-4.24 |
| 2014 | GoF | 2.40-10.33 | 4.58-28.72 | 4.38-19.28 | 0.44-**1.67** | 0.11-0.43 | 1.11-1.94 | 4.21-4.45 |
| May | BP | 1.52-1.91 | 4.47-6.44 | 2.27-25.84 | 0.73-1.14 | 0.20-0.33 | 0.89-1.64 | 3.89-4.31 |
| 5^th^-10^th^ | ArS | **0.70**-2.29 | 3.21-5-78 | 6.61-7.99 | 1.03-1.59 | 0.19-0.33 | 1.67-2.03 | 4.29-4.41 |
| 2015 | GoF | 3.19-12.56 | 5.44-17.70 | **0.28**-5.78 | 0.29-0.47 | 0.15-0.31 | 2.57-4.09 | 3.51-3.68 |
| May | ArS | 1.66-8.90 | 3.39-14.79 | 2.19-6.41 | 0.17-0.25 | 0.08-0.11 | 2.64-4.54 | 3.38-3.56 |
| 4^th^-15^th^ | ÅS | 2.88-4.82 | 4.04-5.50 | 1.58-2.66 | 0.10-0.21 | 0.06-0.25 | 2.62-2.93 | 3.42-3.48 |
|  | BP | 2.88-6.57 | 3.39-13.21 | 2.16-8.57 | 0.17-0.28 | 0.09-0.13 | 1.73-3.22 | 3.21-3.58 |
|  | BS | 6.35-12.91 | 4.19-22.65 | 0.94-4.15 | **0.08**-0.18 | **0.04**-0.13 | 1.58-3.46 | 3.59-3.81 |
|  | Kv, BoB | 1.62-6.79 | **1.01**-4.54 | 2.67-9.95 | 0.10-0.70 | 0.04-0.44 | 1.27**-4.87** | 3.15-3.88 |
| 2016 | GoF | 9.36-**26.07** | 5.64-22.41 | 3.07-13.97 | 0.14-0.28 | 0.09-0.32 | 1.95-2.86 | 3.39-3.53 |
| April | BP | 1.66-11.97 | 2.09-11.02 | 5.91-**21.97** | 0.08-0.25 | 0.05-0.17 | 0.72-2.48 | **2.42**-3.34 |
| 4^th^-15^th^ | ÅS | 7.42-10.16 | 10.42-20.37 | 7.89-15.31 | 0.16-0.19 | 0.10-0.12 | 2.41-3.15 | 3.24-3.41 |

**Table S3.**  Mantel test based of spearman correlations between the taxonomic dissimilarity matrices of bacterial and phytoplankton communities, for each of the sampling cruises. R coefficients are indicated as r value and significance as *p* value (*p* <0.05). N = 122

| Mantel test | r value | p value |
| --- | --- | --- |
| **2013** | 0.754 | 0.0001 |
| **2014** | 0.627 | 0.0001 |
| **2015** | 0.783 | 0.0001 |
| **2016** | 0.784 | 0.0001 |

**Table S4.** Spearman correlations between bacterial production based on thymidine (BPT) and leucine (BPL) incorporation and the relative abundance of main bacterial groups. The correlation is indicated as rho value. N = 122; significant level, *p* < 0.05.

| Spearman  correlation | BPT  rho value | p value | BPL  rho value | p value |
| --- | --- | --- | --- | --- |
| **SAR 11** | -0.636 | < 0.0001 | -0.634 | < 0.0001 |
| **Rhodobacteraceae** | -0.006 | 0.509 | -0.007 | 0.439 |
| **Other Alphaproteobacteria** | -0.671 | < 0.0001 | -0.667 | < 0.0001 |
| **Phycisphaerae** | -0.017 | 0.850 | 0.015 | 0.870 |
| **Acidimicrobiia** | 0.105 | 0.252 | -0.130 | 0.154 |
| **Actinobacteria** | 0.662 | < 0.0001 | 0.471 | < 0.0001 |
| **Betaproteobacteria** | 0.706 | < 0.0001 | 0.653 | < 0.0001 |
| **Gammaproteobacteria** | 0.520 | < 0.0001 | 0.598 | < 0.0001 |
| **Sphingobacteria** | 0.687 | < 0.0001 | 0.570 | < 0.0001 |
| **Flavobacteriia** | 0.625 | < 0.0001 | 0.656 | < 0.0001 |
| **Cytophagia** | 0.698 | < 0.0001 | 0.644 | < 0.0001 |
